# Primary infection with dengue or Zika virus does not affect the severity of heterologous secondary infection in macaques

**DOI:** 10.1101/613802

**Authors:** Meghan E. Breitbach, Christina M. Newman, Dawn M. Dudley, Laurel M. Stewart, Matthew T. Aliota, Michelle R. Koenig, Phoenix M. Shepherd, Keisuke Yamamoto, Chelsea M. Crooks, Ginger Young, Matthew R. Semler, Andrea M. Weiler, Gabrielle L. Barry, Holly Heimsath, Emma L. Mohr, Jens Eichkoff, Wendy Newton, Eric Peterson, Nancy Schultz-Darken, Sallie R. Permar, Hansi Dean, Saverio Capuano, Jorge E. Osorio, Thomas C. Friedrich, David H. O’Connor

## Abstract

Zika virus (ZIKV) and dengue virus (DENV) are genetically and antigenically related flaviviruses that now co-circulate in much of the tropical and subtropical world. The rapid emergence of ZIKV in the Americas in 2015 and 2016, and its recent associations with Guillain-Barré syndrome, birth defects, and fetal loss have led to the hypothesis that DENV infection induces cross-reactive antibodies that influence the severity of secondary ZIKV infections. It has also been proposed that pre-existing ZIKV immunity could affect DENV pathogenesis. We examined outcomes of secondary ZIKV infections in three rhesus and fifteen cynomolgus macaques, as well as secondary DENV-2 infections in three additional rhesus macaques up to a year post-primary ZIKV infection. Although cross-binding antibodies were detected prior to secondary infection for all animals and cross-neutralizing antibodies were detected for some animals, previous DENV or ZIKV infection had no apparent effect on the clinical course of heterotypic secondary infections in these animals. All animals had asymptomatic infections and, when compared to controls, did not have significantly perturbed hematological parameters. Rhesus macaques infected with DENV-2 approximately one year after primary ZIKV infection had higher vRNA loads in plasma when compared with serum vRNA loads from ZIKV-naive animals infected with DENV-2, but a differential effect of sample type could not be ruled out. In cynomolgus macaques, the serotype of primary DENV infection did not affect the outcome of secondary ZIKV infection.

**Author summary:** Pre-existing immunity to one of the four DENV serotypes is known to increase the risk of severe disease upon secondary infection with a different serotype. Due to the antigenic similarities between ZIKV and DENV, it has been proposed that these viruses could interact in a similar fashion. Data from in vitro experiments and murine models suggests that pre-existing immunity to one virus could either enhance or protect against infection with the other. These somewhat contradictory findings highlight the need for immune competent animal models for understanding the role of cross-reactive antibodies in flavivirus pathogenesis. We examined secondary ZIKV or DENV infections in rhesus and cynomolgus macaques that had previously been infected with the other virus. We assessed the outcomes of secondary ZIKV or DENV infections by quantifying vRNA loads, clinical and laboratory parameters, body temperature, and weight for each cohort of animals and compared them with control animals. These comparisons demonstrated that within a year of primary infection, secondary infections with either ZIKV or DENV were similar to primary infections and were not associated with enhancement or reduction in severity of disease based on the outcomes that we assessed.

## Introduction

The spread of Zika virus (ZIKV) from Africa to Asia and the Americas has led to the co-circulation and co-infection of ZIKV with other endemic arboviruses including dengue virus (DENV) (1–4). ZIKV exists as a single serotype while DENV consists of four antigenically similar serotypes (DENV-1-DENV-4) (5, 6). Primary infection with one of the four DENV serotypes typically confers antibody-mediated lifelong protection against reinfection with the same serotype. However, primary DENV infection may protect against, have no effect on, or enhance subsequent infection with a heterotypic serotype (7). In humans, reinfection with a heterotypic DENV serotype is associated with higher viral load and an elevated risk of dengue hemorrhagic fever (DHF) and dengue shock syndrome (DSS) (8). Secondary DENV infections in macaques share some similarities with human infections including a trend toward higher peak DENV viral load (9). In addition, a single rhesus macaque was previously reported as developing clinical responses consistent with dengue fever/DSS in response to secondary DENV-2 infection, including leukocytosis, thrombocytopenia, and elevated hematocrit approximately 5 days after viremia was first detected (9).

The similarity of DENV and ZIKV antigenic epitopes makes it difficult to distinguish between these viruses serologically in people living in DENV/ZIKV endemic areas (10–12), suggesting that pre-existing immunity to either virus might affect the course of infection with the other. Cross-reactive DENV antibodies have been hypothesized as a factor driving the association of ZIKV with Guillain-Barré and adverse pregnancy outcomes (11). Enhancement of ZIKV in the presence of anti-DENV antibodies has been demonstrated both in vitro and in vivo, in particular in murine models (13–16). Experimental infections in mice have shown enhancement of, and protection from, ZIKV infection in the context of DENV immunity (15, 17–19). In humans to date, enhancement of ZIKV infection by pre-existing DENV immunity has not been observed (20). In fact, a recent study in Salvador, Brazil showed an association between high anti-DENV total IgG titers and a decreased risk of ZIKV infection and symptoms (21). Likewise, clinical data collected from pregnant women in Rio de Janeiro also suggest that there may be no association between the presence of DENV antibodies and ZIKV-associated pregnancy outcomes; however, DENV seroprevalence in the study population was >88% (22).

Macaque monkeys are commonly used in biomedical research as models for human diseases (23). One key advantage of macaques over other animal models for studying vector-borne flaviviruses is their susceptibility to infection without requiring immunological manipulation. Primary ZIKV infections in macaques range from subclinical infections to mild fever, conjunctivitis, and rash (24–30). In primary DENV infection in macaques, severe disease, including hemorrhage, has only been induced using a high dose intravenous inoculation (31). On the other hand, infections with doses designed to mimic mosquito-borne transmission of DENV are clinically inapparent, necessitating the use of secondary clinical and laboratory parameters such as viral RNA (vRNA) loads, complete blood count (CBC) tests, and serum chemistry panels to study disease enhancement (31). Multiple nonhuman primate studies utilizing a variety of DENV serotypes and ZIKV strains have been done over variable periods of time following primary infection to investigate the impact of primary DENV-1, DENV-2, and DENV-4 infection on secondary ZIKV infection; however, none to date have documented ZIKV enhancement (24–30). Notably, no studies have looked at prior infection with DENV-3. Our study more than doubles the total number of macaques used to study DENV/ZIKV interactions and incorporates data from all four DENV serotypes.

Here we examined whether laboratory markers of clinical illness and vRNA loads are enhanced during secondary heterologous flavivirus infections in macaques. We also differentiate changes in clinical and laboratory parameters that are due to sedation, stress, and frequent venipuncture on animals from the impact of ZIKV infection, which were not controlled in previous studies. This work is timely given the efforts to develop an effective vaccine for ZIKV, as well as the introduction of a tetravalent DENV vaccine candidate, and other DENV vaccines in areas where ZIKV and DENV are now co-endemic (32, 33). This study adds new, more comprehensive and controlled information about the impact of prior DENV infection on secondary ZIKV disease in the macaque model.

## Results

### Primary DENV-3 infection does not enhance secondary ZIKV infection in rhesus macaques

#### ZIKV plasma vRNA load in DENV-3 immune animals is similar to ZIKV plasma vRNA load in naive animals

No studies to date have examined the impact of primary DENV-3 exposure on secondary ZIKV infection in a rhesus macaque model. Previous studies have also not included mock ZIKV infections to control for the possibility that changes in clinical and laboratory parameters are due to stressful, frequent animal handling and venipuncture.

Three Indian rhesus macaques (*Macaca mulatta*) were inoculated subcutaneously (SC) with Asian-lineage ZIKV (Zika virus/H.sapiens-tc/FRA/2013/FrenchPolynesia-01-v1c1; ZIKV-FP) approximately 1 year (850858 and 489988) or 0.4 years (321142) after infection with DENV-3 (dengue virus/H.sapiens-tc/IDN/1978/Sleman/78) (Table 1, cohort A and **S1 Table**). A sampling timeline for viral RNA (vRNA) load and all other parameters is shown in Fig 1. Plasma ZIKV vRNA was detectable by 1 day post-infection (dpi) and cleared by 8 dpi for 850585 and 321142 and by 10 dpi for 489988 (Fig 2). Peak ZIKV RNA loads ranged from 3.5×10^5^ to 3.0×10^6^ vRNA copies/mL plasma at 3 or 4 days post-ZIKV exposure (Fig 2). The magnitude of plasma ZIKV vRNA burden did not differ between cohort A and ZIKV control animals (Mann-Whitney U-test, W = 5, p = 0.57) that were infected with the same dose, route, and strain of ZIKV in previous studies (Table 1 **and** Fig 2) (27, 34). Likewise, we did not observe a significant difference in ZIKV vRNA load overall between cohort A animals and ZIKV control animals based on comparison of the area under the curve (AUC) for each group (Student’s t-test, t (5.96) = −1.89, p = 0.11) (Fig 2). One animal in cohort A (489988) and one ZIKV control animal (448436) had undetectable vRNA loads at 8 dpi, but were detectable again at 9 dpi. All plasma vRNA loads resolved by 14 dpi (Fig 2).

**Table 1.** Brief summary of each cohort including viruses used to infect animals, animal IDs, route of virus exposure, and animal species.

| Cohort | 1° Infection | 2° Infection | Animal ID | Exposure Route | Species |
| --- | --- | --- | --- | --- | --- |
| A | DENV-3 | ZIKV-FP | 850585<br>489988<br>321142 | Subcutaneous | Indian Rhesus Macaque |
| B | ZIKV-FP | DENV-2 | 912116<br>393422<br>826226 | Subcutaneous | Indian Rhesus Macaque |
| C-1 | DENV-1 | ZIKV-PR | 940262<br>289427 | Mosquito Bite | Mauritian Cynomolgus Macaque |
|  |  |  | 941637 | Subcutaneous |  |
| C-2 | DENV-2 | ZIKV-PR | 638166<br>346817<br>658991<br>714622 | Mosquito Bite | Mauritian Cynomolgus Macaque |
| C-3 | DENV-3 | ZIKV-PR | 644369<br>865011<br>359807 | Mosquito Bite | Mauritian Cynomolgus Macaque |
|  |  |  | 753662 | Subcutaneous |  |
| C-4 | DENV-4 | ZIKV-PR | 523664<br>820832<br>456891<br>624311 | Mosquito Bite | Mauritian Cynomolgus Macaque |
| negative control | PBS | NA | 774011<br>829256<br>875914 | Subcutaneous | Indian Rhesus Macaque |
| DENV control | DENV-2 | NA | rh2484<br>rh2486<br>rh2495<br>rh2499<br>rh2503 | Subcutaneous | Indian Rhesus Macaque |
| ZIKV control | ZIKV-FP | NA | 448436<br>756591<br>861138<br>411359<br>912116 | Subcutaneous | Indian Rhesus Macaque |

**Fig 1.**
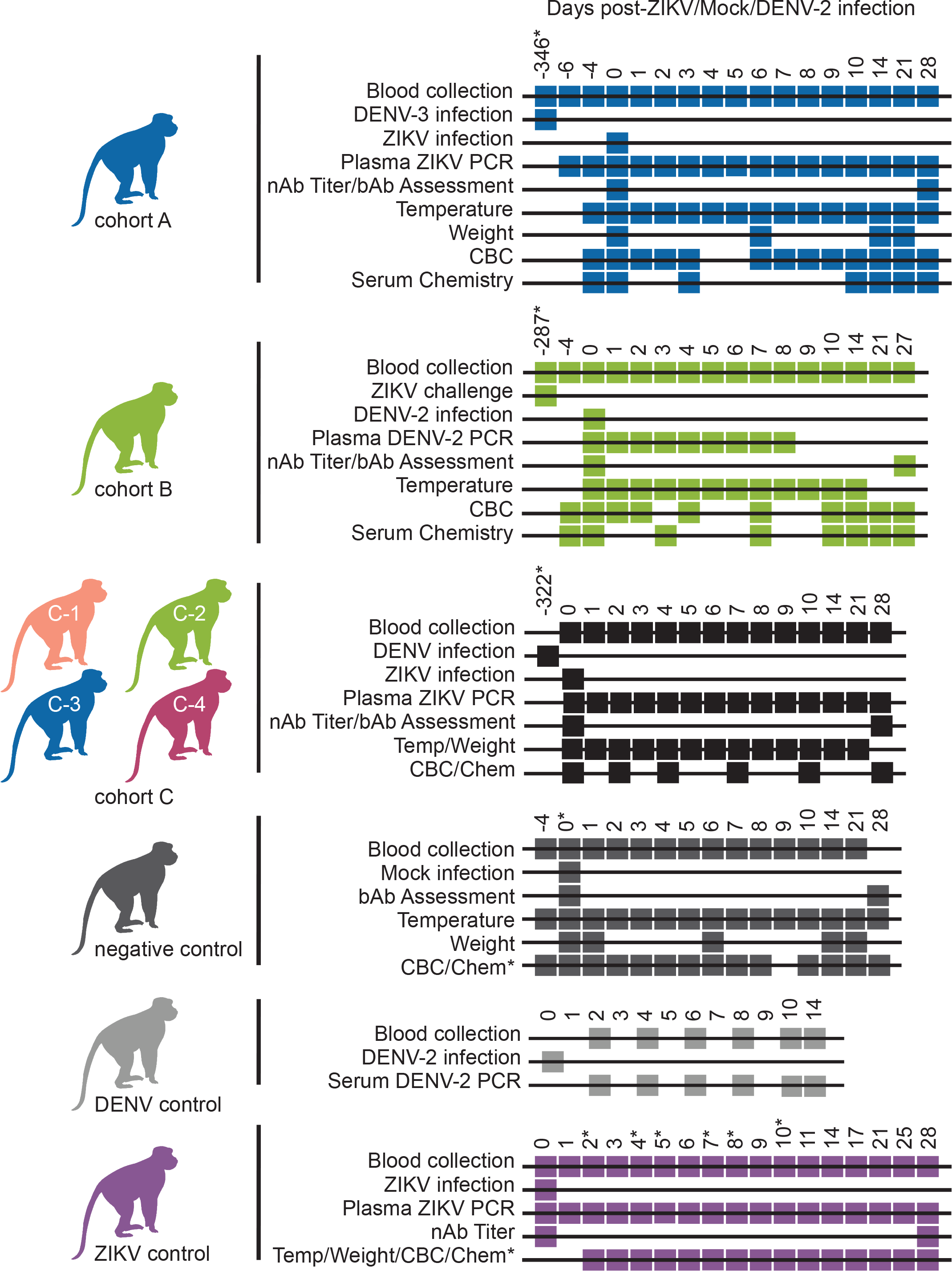
Sampling timeline for each cohort. Sampling and infection schema for all groups of animals presented in this study. Days where an inoculation was performed or samples were collected to run a test are annotated with a box at the time point relative to either the primary viral infection in the DENV control and ZIKV control groups or relative to the secondary infection in cohorts A-C. Secondary infections in cohorts A and C were with ZIKV while secondary infection in cohort B was with DENV. Asterisks denote when the sampling for an animal/animals differed from the Cohort and is as follows: For cohort A, 321142 was infected with ZIKV 164 days after primary exposure to DENV-3. Cohort B animals were challenged with ZIKV twice before DENV infection. The cohort B timeline is shown relative to secondary ZIKV challenge as opposed to primary ZIKV infection, which occurred 357 days prior to DENV-2 infection. For cohort C, animals were infected with ZIKV in two groups and the timeline begins relative to ZIKV infection of the first group of animals. Animals in the second group were infected with DENV 351 days prior to ZIKV infection. Cohort C animals 752662 and 941637 were infected with DENV 387 days prior to SC inoculation with ZIKV. Serum chemistry panels for negative control animals were analyzed on days −4, 0, 3, 10, 14, 21, and 28, except for 774011 who was not sampled on day 0. Day 5 post-infection CBC tests were not included for negative control animals due to incorrect blood collection. Serum chemistry panels were analyzed for ZIKV control animal 411359 on days 0, 1, 2, 3, 4, 6, and 14 while for ZIKV control animal 912116 they were analyzed on days −6, 2, 5, and 11 relative to ZIKV infection. CBC tests were analyzed for ZIKV control animals on days −4, 0, 1, 2, 3, 4, 6, 7, 8, 9, 14, 21, and 28, except for 912116 for whom these data were not collected on days 2, 4, 7, 8, and 10 relative to ZIKV infection. For animals without day −4 samples, day 0 samples were considered baseline samples and vice versa.

**Fig 2.**
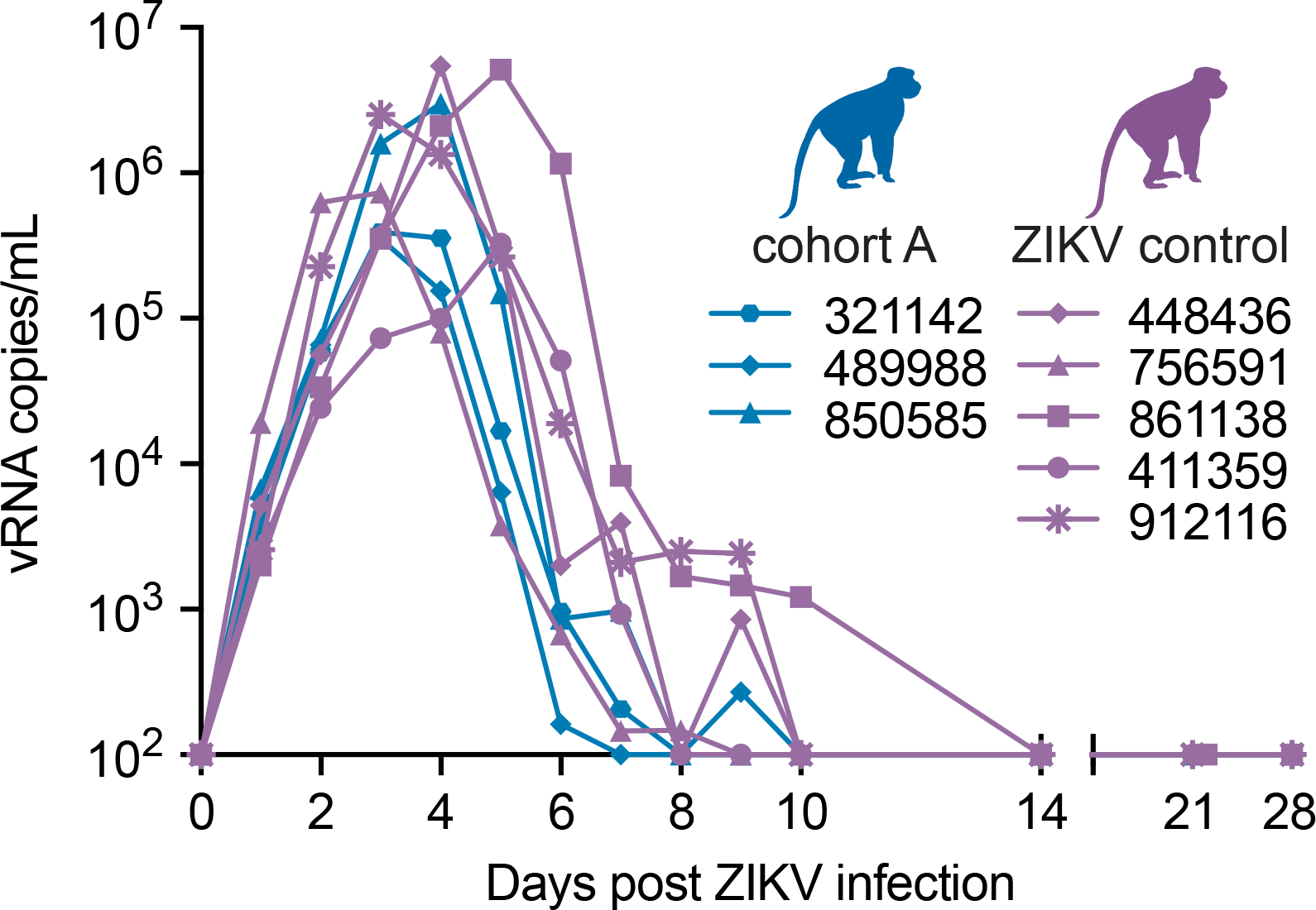
Cohort A ZIKV viral loads do not differ from ZIKV control animals. Longitudinal viral RNA copies/mL of plasma is plotted after ZIKV infection for cohort A and ZIKV control groups (see Legend). The viral load graph starts at the limit of detection of the ZIKV viral load assay, which is 100 copies vRNA/ml plasma.

#### Prior DENV-3 exposure does not generate cross-reactive antibody responses to ZIKV

Similar to sequential heterotypic DENV infections, the effect of prior DENV exposure on ZIKV infection is likely determined by the nature of the antibody response at the time of secondary infection. Neutralizing antibodies (nAbs) are associated with protection while binding antibodies (bAbs) are associated with enhancement (35, 36). To evaluate cross-neutralizing antibody responses against ZIKV from prior DENV-3 infection, both ZIKV and DENV nAb titers were measured at 0 and 28 days post-ZIKV infection by plaque reduction neutralization test (PRNT). On the day of ZIKV infection, DENV-3 immune sera did not cross-neutralize ZIKV (Fig 3A). Neutralizing antibody titers above 1:10 against DENV-3 present just prior to ZIKV infection serve as confirmation of DENV-3 immunity in these animals (Fig 3B). By 28 days post-ZIKV infection, serum neutralized both ZIKV and DENV-3 (PRNT_50_ titers > 1:1000). Interestingly, ZIKV infection boosted DENV-3 nAb titers to PRNT_50_ titers ≥ 1:1219, indicating that nAbs generated in response to ZIKV infection cross-neutralize DENV-3 (Fig 3B).

**Fig 3.**
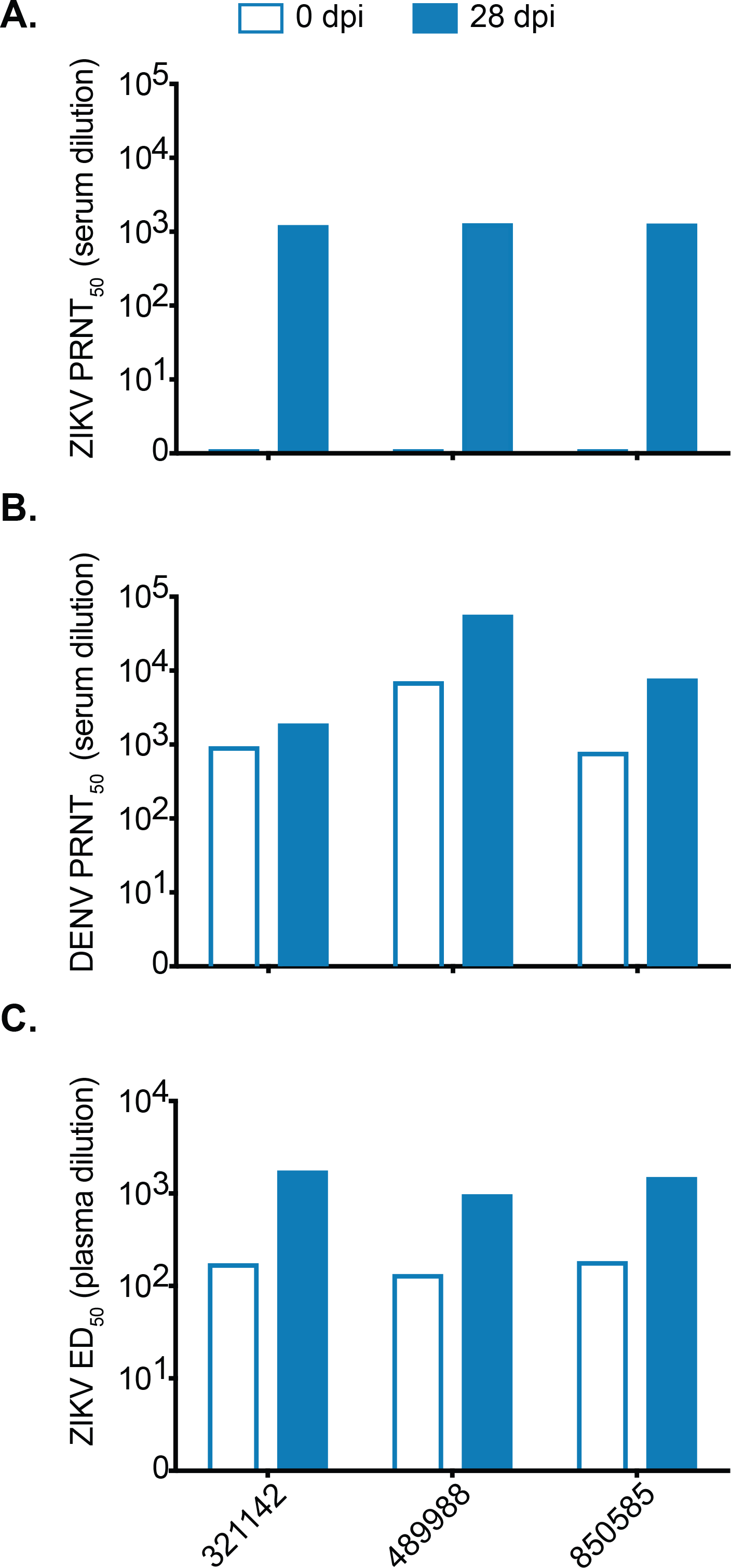
Prior DENV infection generates cross-binding but not cross-neutralizing ZIKV antibodies. (A) ZIKV-specific neutralizing antibody titer of cohort A animals before (open bars) and 28 days after infection (filled bars) with ZIKV determined by a 50% plaque neutralization test. (B) DENV-specific neutralizing antibody titer of cohort A animals before and 28 days after infection with ZIKV. (C) Whole virion ZIKV-specific binding antibody levels of cohort A animals expressed as log_10_ 50% effective dilution of binding in an ELISA assay before and 28 days after ZIKV infection.

Antibodies that bind rather than neutralize, are associated with ADE in heterotypic DENV infections (35–37). Therefore, we also used a ZIKV whole-virion binding ELISA to detect all antibodies that bind to ZIKV, regardless of neutralizing capacity, in the plasma of cohort A animals before and after ZIKV infection. All three cohort A animals had detectable ZIKV-bAb immediately before ZIKV infection with the log_10_ 50% effective dilution (ED_50_) at > 1:100 (Fig 3C). Since these animals did not have detectable nAbs, it is reasonable to surmise that the bAbs detected at baseline are not neutralizing. At 28 days post-ZIKV infection, the ED_50_ for cohort A animals increased by approximately 1 log_10_, indicating a boost of ZIKV bAbs after ZIKV infection.

#### Clinical and laboratory parameters do not suggest enhancement of ZIKV pathogenesis in DENV-3 immune rhesus macaques

A small number of severe human clinical cases have reported thrombocytopenia, anemia, and hemorrhagic manifestations detected by serum chemistry and complete blood count (CBC) tests (38). To investigate ZIKV-associated disease in the absence of outward clinical signs, serum chemistry panels, CBCs, body temperature, and body weight data were evaluated **(S1 and S2 Figs)**. Chemistry panel values were normalized to baseline by calculating the fold change from baseline in each parameter for each animal and were analyzed using a linear mixed effects model. Cohort A animals showed no significant differences overall in any parameter when compared with negative control animals **(S1 Fig,** see **https://go.wisc.edu/1u3nu1** for full list of p-values for pairwise comparisons). This indicates that fluctuations from baseline in cohort A may be attributed to the animal handling and sampling stresses also received by the negative control animals, rather than due to secondary ZIKV infection as shown in **S1 and S2 Figs**. Interestingly, the ZIKV control group had lower serum levels of both aspartate aminotransferase (AST) and lactate dehydrogenase (LDH) when compared to both cohort A and negative control animals **(S1 Fig E, F)**. These differences may not be clinically meaningful due to the small number of ZIKV control animals (n = 2) and the significant individual variation, likely in response to the stress of sedation and manipulation during the study. Consistent with daily blood sampling through day 10, both hemoglobin (HB) and hematocrit (HCT) decreased from baseline in all groups until 10 dpi **(S2 Fig A, B)**. Platelet and WBC counts showed fluctuations from baseline at the individual level; however, like the serum chemistry panel results, the consistency of these findings with those of the negative control animals suggests that the observed fluctuations in CBC parameters were not the result of ZIKV infection **(S2 Fig C, D)**. CBC test parameters were normalized by calculating the fold change from baseline and then analyzed using a linear mixed effects model. Pairwise comparisons between cohort A, the ZIKV control group, and negative control group showed no significant difference between groups for any CBC test parameter **(S2 Fig, https://go.wisc.edu/1u3nu1).** Likewise, no significant difference in body temperature (t(7) = −1.05, p = 0.32) nor body weight (t(42) = 1.36, p = 0.18) were observed **(S2 Fig E, F)**. No animals exhibited an elevated body temperature above the normal range for rhesus macaques based on WNPRC reference ranges and overall weight changes were minimal **(S2 Fig E, F)**.

#### Minor cytokine and chemokine fluctuations associated with secondary ZIKV infections in DENV-3 immune rhesus macaques

In humans, enhanced DENV infection is accompanied by increases in pro-inflammatory and vasoactive cytokines that contribute to vascular leakage and hemorrhagic fever (39–42). To understand whether perturbations in cytokines were detectable in our animals despite no clinical signs of disease, we longitudinally compared the cytokine and chemokine profiles of cohort A animals to ZIKV control and negative control animals with a 23-plex primate panel using the Luminex platform throughout the 28-day study. This panel detects multiple cytokines associated with DENV infection and enhanced DENV disease including MCP-1, TNF-ɑ, IL-8, IL-15, IFN-Ɣ, IL-1ra, and IL-4 (42–45). Most parameters remained below the limit of detection for the assay throughout the study, except for monocyte chemoattractant protein 1 (MCP-1), sCD40L, IL-1ra, IL-2, IL-8, and IL-15. Cohort A animals showed no significant difference in levels of sCD40L, IL-8, or IL-2 when compared with negative control or ZIKV control animals **(S3 Fig A, B, C)**. Both IL-1ra and IL-15 significantly increased in plasma of both cohort A and ZIKV controls compared with negative control animals based on MANOVA with pairwise post-hoc comparisons (Bonferroni adjusted p =4.3×10^−7^and 1.3×10^−6^ respectively for IL-1ra and p = 0.0007 and p = 1.3×10^−7^ respectively for IL-15) **(S3 Fig D, E)**. In addition, IL-15 trended lower for ZIKV control animals when compared with cohort A (p = 0.046). MCP-1 levels were highest in ZIKV control animals (compared with cohort A: p = 0.02 and negative controls: p < 0.0001) followed by cohort A (compared with negative controls: p = 0.0001) based on repeated measures ANOVA with post-hoc analysis using Tukey’s HSD **(S3 Fig F)**. It should be noted that animal 850585 had a small scratch noted on his nose at 9 dpi, likely contributing to elevated levels of IL-2, sCD40L and IL-1ra starting at this time-point. Omitting this animal did not alter the statistical interpretation of these data and therefore he was included. Altogether, ZIKV infection led to an increase in IL-1ra, IL-15, and MCP-1 in all infected animals relative to negative control animals, but only MCP-1 differed between cohort A and ZIKV control animals.

### Prior ZIKV infection does not result in clinical disease in rhesus macaques during secondary DENV-2 infection

#### Peak DENV-2 plasma vRNA load was increased in ZIKV immune rhesus macaques when compared with peak DENV-2 serum vRNA load of historical controls

To explore whether previous ZIKV exposure impacts subsequent DENV disease, we infected three Indian rhesus macaques, exposed twice to ZIKV-FP in previous studies, with DENV-2 (dengue virus/H.sapiens-tc/NGU/1944/NGC-00982-p17c2; New Guinea C) (cohort B) (Table 1, **S1 Table and** Fig 1). Cohort B animals had detectable DENV RNA loads in plasma by 1 (393422, 912116) and 2 dpi (826226) (Fig 4). Peak DENV plasma vRNA loads occurred at 6 dpi for all animals and ranged between 1.2×10^6^−2.3×10^6^ vRNA copies/mL (Fig 4). Five ZIKV-naive rhesus macaques from a prior study were used as DENV control animals and were SC-inoculated with DENV-2 (Fig 4). Serum, but not plasma samples, were collected from DENV control animals every other day 4 from day 0 through day 14 post-infection. From these sample time-points, peak serum vRNA loads occurred on day 6 post-infection and ranged from 3.9×10^4^−1.0×10^5^ vRNA copies/mL. Unfortunately, since DENV-2 vRNA data were collected from two different studies, all cohort B vRNA loads were quantified from plasma whereas all DENV control vRNA loads were quantified from serum. When comparing vRNA loads between these cohorts and sample types, cohort B animals had higher peak vRNA loads than DENV control animals at 6 dpi (Student’s t-test, t(2) = − 4.53, p = 0.045), likely because serum vRNA loads are often lower than temporally matched plasma vRNA loads for both ZIKV and DENV **(S4 Fig)**. It is possible that the difference we observed for 6 dpi vRNA loads between plasma from cohort B animals, and serum from DENV control animals, is due to comparing plasma versus serum rather than prior ZIKV exposure.

**Fig 4.**
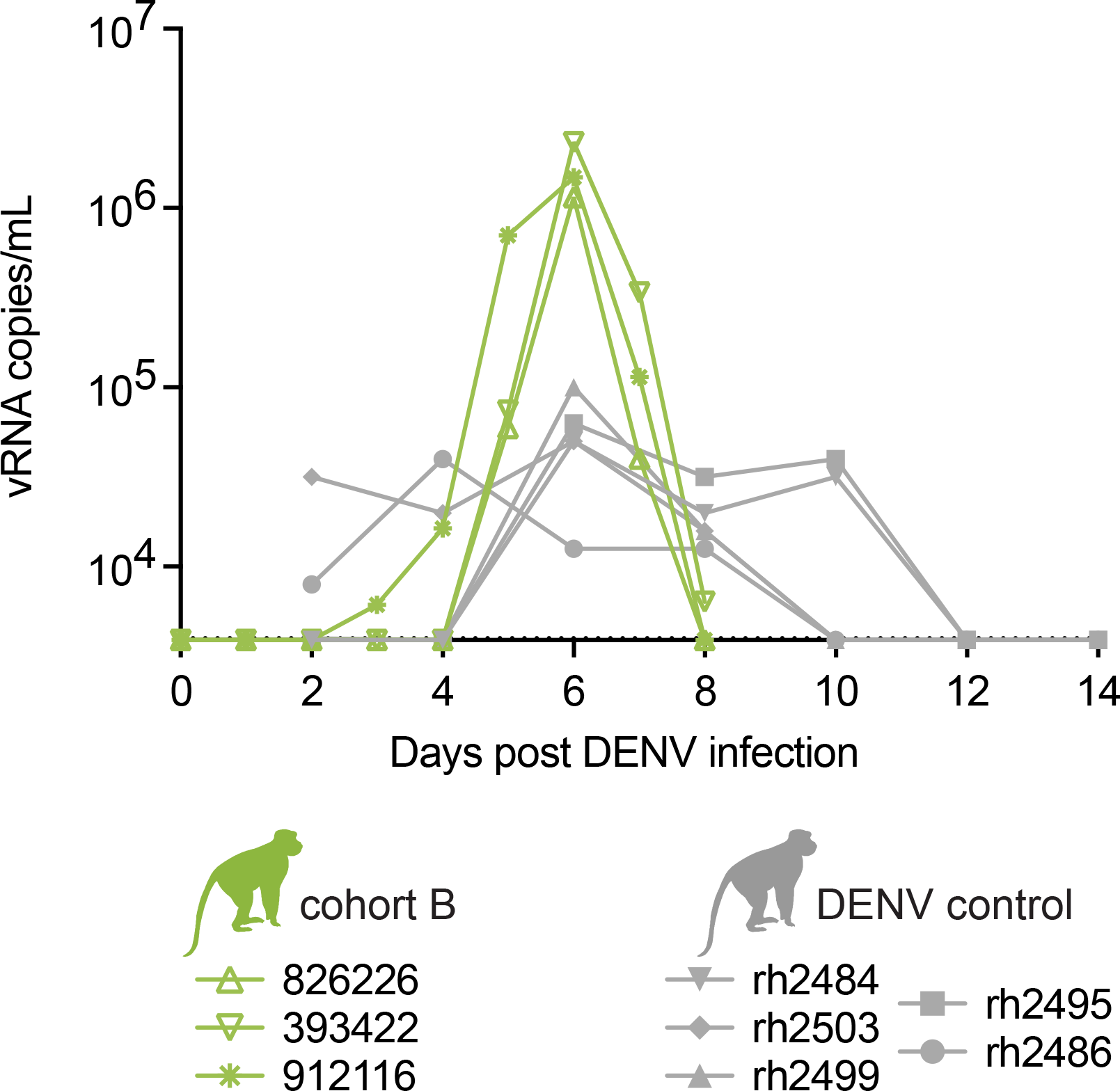
Cohort B DENV viral loads from plasma are higher than DENV control viral loads from serum. Longitudinal viral RNA copies/mL of cohort B animal plasma and DENV control animal serum. The viral load graph starts at the limit of detection of the DENV viral load assay, which is 1 × 10^3.6^ copies vRNA/ml plasma or serum.

#### Prior ZIKV exposure generates cross-reactive antibody responses to DENV

To quantify ZIKV- and DENV-specific nAb titers in cohort B animals at the time of secondary DENV infection, serum was tested immediately before and at 28 days post-DENV infection using both ZIKV and DENV PRNT_50_. Before DENV infection (0 dpi), ZIKV immune sera from all cohort B animals cross-neutralized DENV-2 at low levels (PRNT_50_ titers ≤ 1:30) and potently neutralized ZIKV (PRNT_50_ titers 1:1294-1:3954) (Fig 5A, B). At 28 days after DENV-2 infection, DENV-2 PRNT_50_ titers increased to 1:176-1:1570 while ZIKV PRNT_50_ titers also increased, but by less than ten-fold (Fig 5A, B).

**Fig 5.**
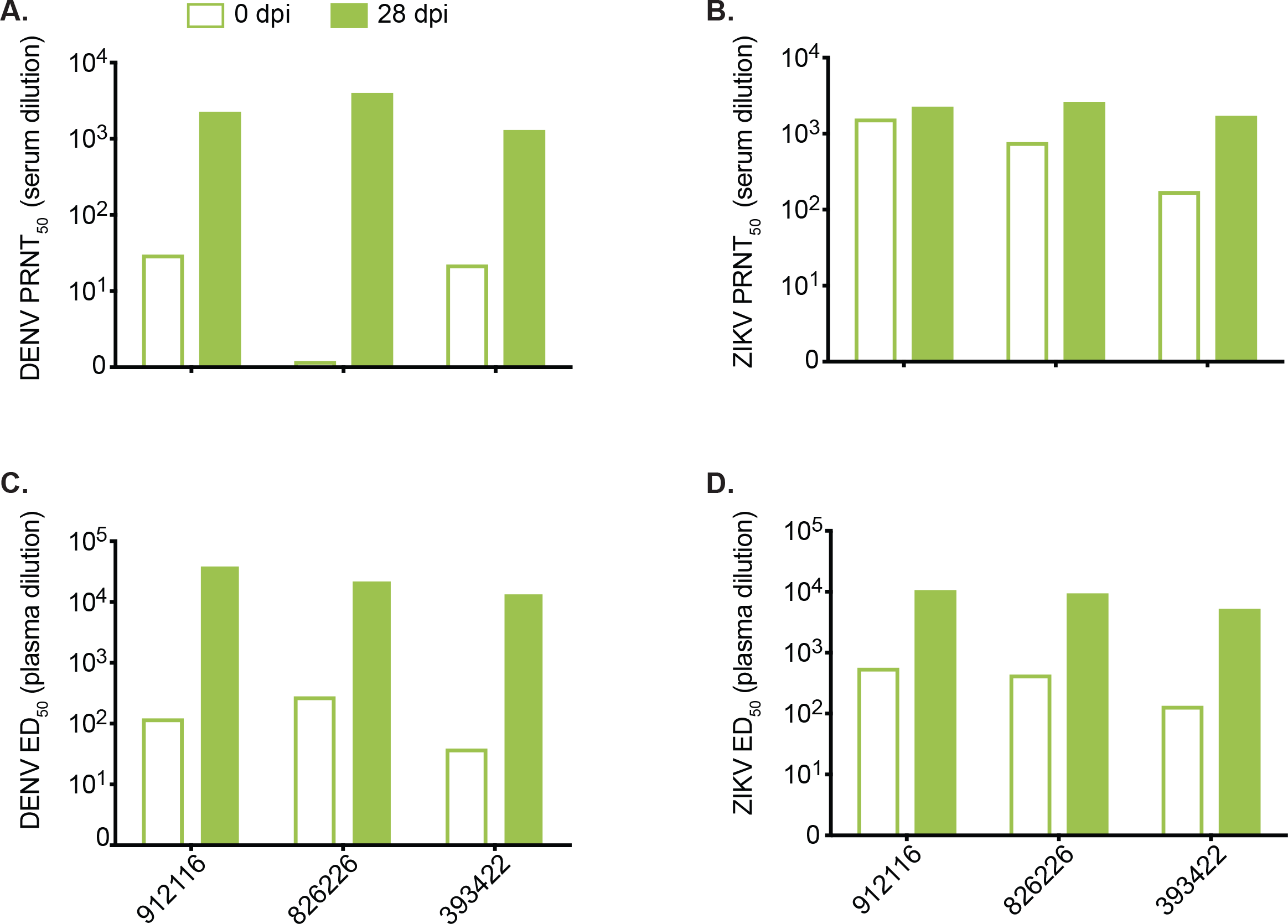
Prior ZIKV infection generates cross-binding and cross-neutralizing DENV antibodies that are boosted after DENV infection. (A) DENV-specific neutralizing antibody titer of cohort B animals before (open bars) and 28 days after DENV infection (filled bars) determined by a 50% plaque neutralization test. (B) ZIKV-specific neutralizing antibody titer of cohort B animals before and 28 days after infection with DENV determined by a 50% plaque neutralization test. (C) Whole virion DENV-specific binding antibody levels of cohort B animals expressed as log_10_ 50% effective dilution of binding in an ELISA assay before and 28 days after ZIKV infection. (D) Whole virion ZIKV-specific binding antibody levels of cohort B animals before and 28 days after ZIKV infection.

The presence of bAbs to ZIKV-PR and DENV-2 was assessed using whole-virion binding ELISA on plasma collected at 0 and 28 days post-DENV-2 infection. Plasma from all cohort B animals contained low levels of cross-reactive bAbs to DENV-2 before infection with DENV-2 (PRNT_50_ titers > 1:100) which increased by day 28 post-DENV-2 infection (Fig 5C). Anti-ZIKV bAbs were present at 0 days post-DENV infection and interestingly increased after DENV-2 exposure (Fig 5D). In stark contrast to cohort A results, these results indicate cross-reactivity of nAbs and bAbs elicited by ZIKV infection against DENV prior to DENV infection, as well as increased cross-reactive antibodies to both ZIKV and DENV after DENV infection (Fig 5).

#### Clinical and laboratory parameters do not suggest enhancement of DENV-2 pathogenesis in ZIKV immune rhesus macaques

Primary DENV infections in macaques can be associated with perturbations in WBC counts and other blood homeostatic parameters such as HCT and liver enzymes (ALT and AST) (46). Clinical and laboratory parameters associated with DENV disease were assessed over time using serum chemistry panels, CBC tests, and body temperature, and were compared between cohort B and negative control animals. These data were not available for DENV control animals. Serum chemistry parameter values were normalized by calculating the fold change from baseline and then compared using the AUC for each parameter and group of animals. Overall, cohort B animals showed a decrease in serum ALP relative to baseline when compared with negative control animals (p = 0.0034) **(S5 Fig A)**. However, raw serum ALP baseline values were elevated for cohort B animals compared with both the WNPRC reference range and negative control animals prior to DENV infection, remained elevated even after the apparent decrease after DENV infection, and were not considered to be clinically relevant. Other chemistry panel and CBC test parameters did not differ significantly between cohort B and negative control animals **(S5 Fig B-F, S6 Fig, https://go.wisc.edu/1u3nu1)**. Body temperatures also did not fluctuate significantly over the study period for either cohort B or negative control animals (t(4) = 0.38, p = 0.72) and all stayed within WNPRC reference range for rhesus macaques (36-40°C) **(S6 Fig)**.

### Primary infection with any of the four DENV serotypes does not enhance secondary ZIKV infection in Mauritian cynomolgus macaques

#### Peak ZIKV RNA load and duration of ZIKV vRNA detection in animals with sequential DENV then ZIKV challenge

To directly compare the impact of different DENV serotypes on ZIKV disease, four groups of three or four Mauritian cynomolgus macaques (MCM, *Macaca fascicularis*), each previously infected SC with a single DENV serotype: dengue virus/H.sapiens-tc/NRU/1974/WP74 (hereafter DENV-1), dengue virus/H.sapiens-tc/NGU/1944/NGC-00982–p17c (hereafter DENV-2), dengue virus/H.sapiens—tc/IDN/1978/Sleman/78 (hereafter DENV-3), or dengue virus/H.sapiens-tc/IDN/1978/1228 (hereafter DENV-4) (annotated as cohort C-1 through C-4 respectively), were exposed to a Puerto Rican ZIKV isolate (ZIKV-PR) approximately one year after exposure to DENV (Table 1, Fig 1, **and S1 Table)**. DENV vRNA loads and PRNT_50_ titers from the primary DENV infections are shown in **S7 Fig**. MCM were infected with ZIKV via *Aedes aegypti* mosquito bite as described previously (47). Mosquito transmitted infections were used to better represent a natural mode of ZIKV transmission **(S1 Table)**. The dose of ZIKV inoculated by a single mosquito was estimated by saliva plaque assay and ranged from 10^1.5^ to 10^3.3^ PFU **(S8 Fig and S1 Table)**. Single animals from cohorts C-1 (941637) and C-3 (753662) were not successfully infected with ZIKV via mosquito bite and were subsequently SC-inoculated with ZIKV-PR to ensure the number of animals in each serotype exposure group remained as consistent as possible.

Peak ZIKV vRNA loads for MCM infected via mosquito bite occurred between 4 and 8 dpi and ranged from 3.0×10^3^ to 6.4×10^5^ vRNA copies/mL plasma (mean = 8.3×10^4^ vRNA copies/mL) (Fig 6). Peak vRNA loads for SC-inoculated animals (941637 and 753662) occurred on 2 (2.3×10^4^ vRNA copies/mL) and 3 dpi (5.3×10^3^ vRNA copies/mL) respectively (Fig 6A, C). The timing of peak vRNA loads for all animals was consistent with previously published data on mosquito-transmitted and SC ZIKV infection of DENV-naive rhesus monkeys (47). No significant differences in overall ZIKV RNA loads (F(3, 11) = 0.922, p = 0.46) nor peak vRNA loads (F(3, 11) = 0.72, p = 0.56) between DENV serotype exposure groups were observed (AUC compared by ANOVA, Fig 6). The two SC-inoculated animals (753662 and 941637) were excluded from the AUC-based vRNA load analyses with their serotype groups because the AUC values for their longitudinal vRNA loads were significantly lower than those of mosquito-infected animals (Student’s t-test, t = 4.7, df = 4.9, p = 0.0053). For mosquito infected animals, we also examined whether the number of days of ZIKV plasma vRNA detection differed by prior DENV serotype exposure using ANOVA and found no significant difference in the number of days of detection by qRT-PCR between prior exposures (F(3, 11) = 0.27, p = 0.85).

**Fig. 6.**
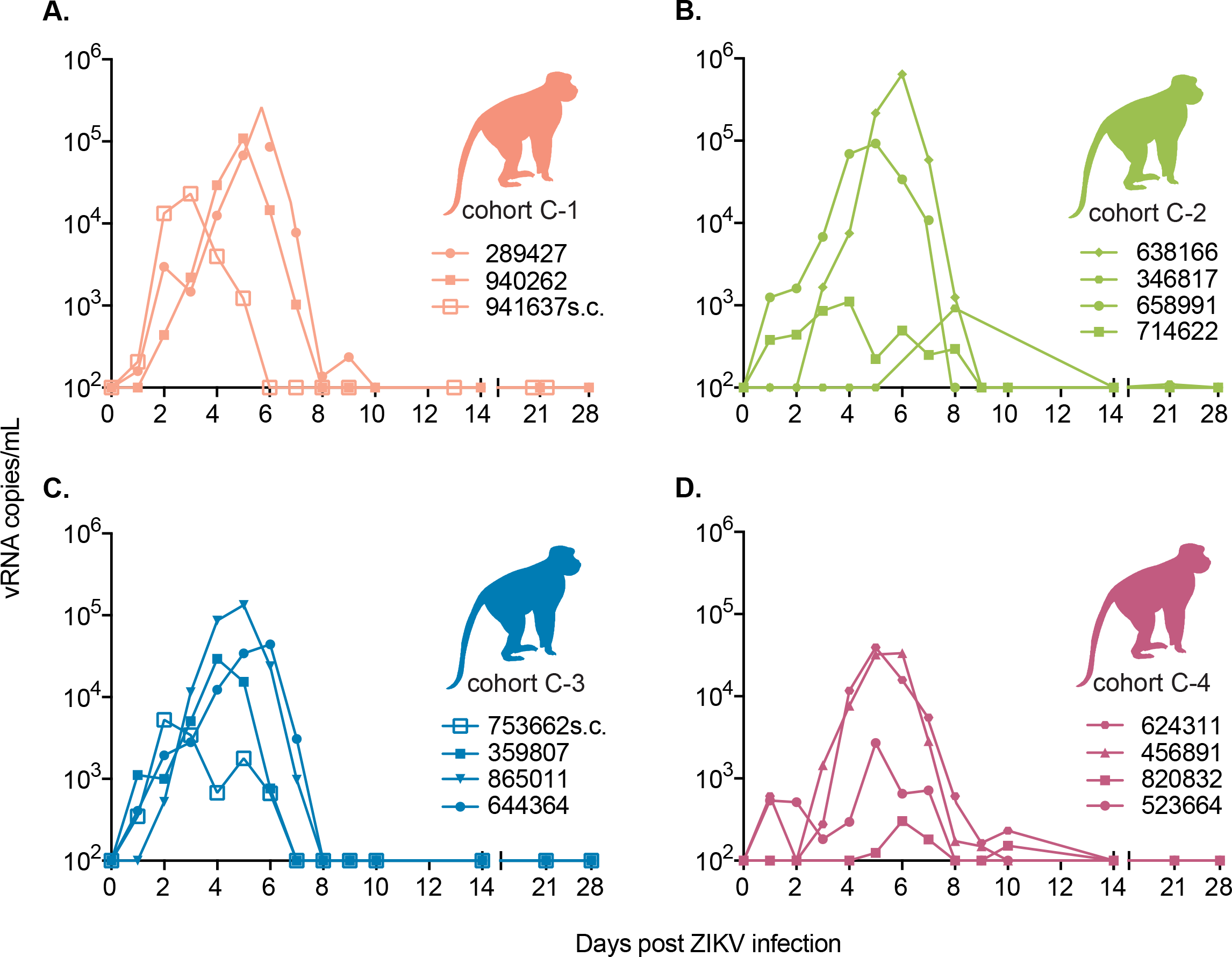
Cohort C ZIKV viral loads do not differ between animals with prior exposure to different DENV serotypes. ZIKV viral RNA copies/ml of plasma over time for animals with prior exposure to DENV. Subcutaneously inoculated animals after failed mosquito bite inoculation are annotated with “SC” after the animal ID and viral loads are presented post-subcutaneous infection. (A) Cohort C-1 animals with prior exposure to DENV-1 (B) Cohort C-2 animals with prior exposure to DENV-2. (C) Cohort C-3 animals with prior exposure to DENV-3. (D) Cohort C-4 animals with prior exposure to DENV-4. The viral load graph starts at the limit of detection of the ZIKV viral load assay, which is 100 copies vRNA/ml plasma.

#### ZIKV cross-neutralizing and binding antibodies in MCM previously exposed to DENV

Serum collected prior to ZIKV infection from animals previously infected with 359 DENV-1 (cohort C-1) showed some cross-neutralization of ZIKV (PRNT_50_ ≤ 1:94), while serum from animals previously infected with either DENV-2, DENV-3, or DENV-4 did not cross-neutralize ZIKV by PRNT_50_ assay (titer range = 1:1.2 - 1:4) (Fig 7A–D). Despite cross-neutralization in animals previously infected by DENV-1, all animals had detectable vRNA in their plasma after exposure to ZIKV-PR following mosquito feeding or SC inoculation with a normal duration of ZIKV viremia lasting 6-10 days for most animals in each group. By 28 or 29 days post ZIKV infection, serum from all cohort C animals potently neutralized ZIKV (PRNT_50_ range: 1:776 - 1:6310) (Fig 7A–D).

We utilized a ZIKV-PR whole-virion binding ELISA to examine whether prior exposure to DENV in MCM resulted in antibodies that bound ZIKV on day 0, prior to ZIKV infection. We found that DENV infection with any serotype resulted in cross-reactive bAbs still detectable approximately one year post-exposure to DENV (Fig 7E **and** Fig 1). As expected, all animals showed an increase in binding based on ED_50_ at 28 days post-ZIKV infection.

**Fig 7.**
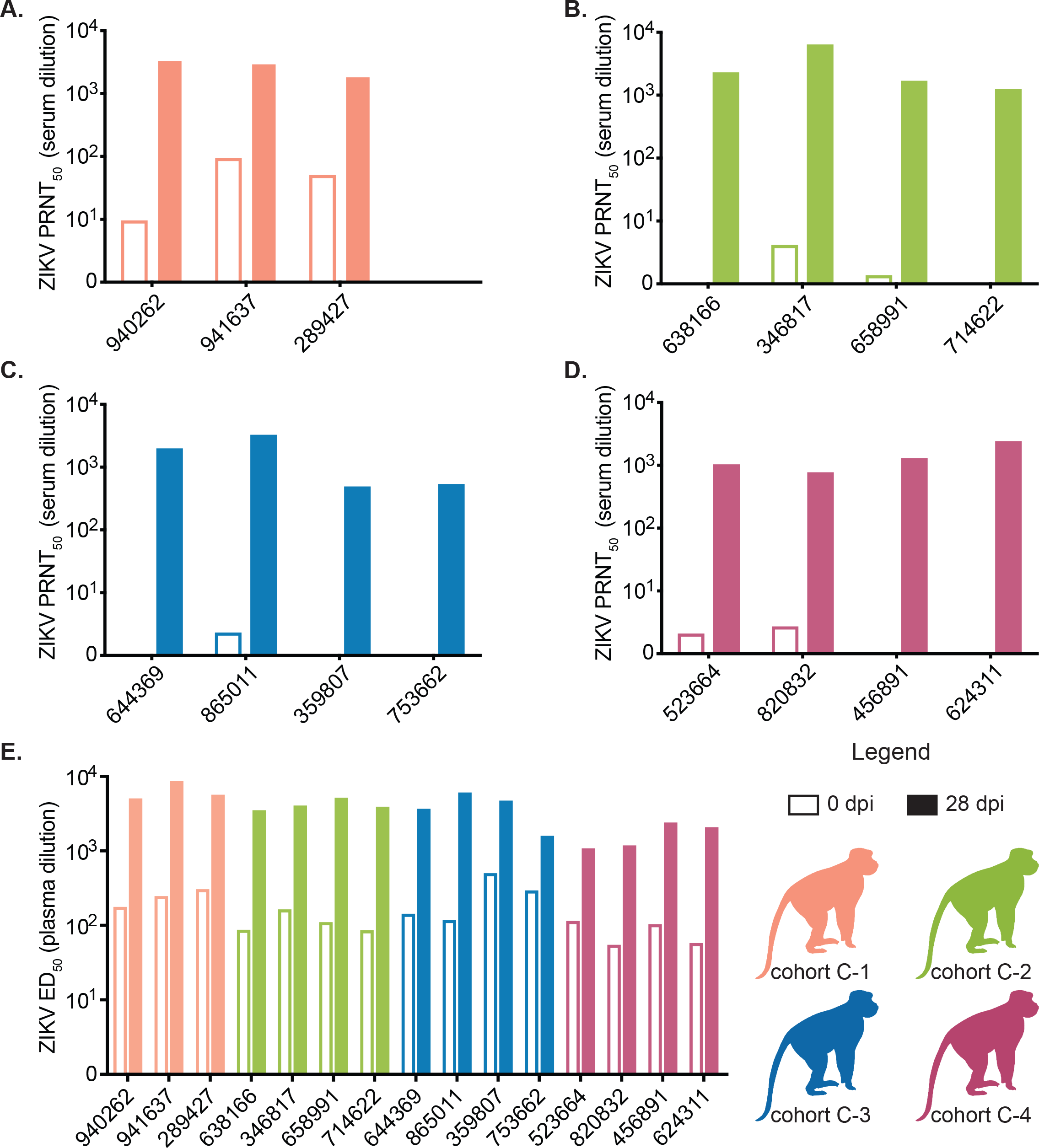
Prior DENV-1 infection generates cross-neutralizing antibodies while all serotypes generate cross-binding antibodies. ZIKV-specific neutralizing antibody titers determined by a 50% plaque neutralization test of cohort C animals before (open bars) and 28 days after ZIKV infection (filled bars). See legend for colors representing each cohort. (A) Antibody titers from cohort C-1. (B) Antibody titers from cohort C-2. (C) Antibody titers from cohort C-3. (D) Antibody titers from cohort C-4. (E) Whole virion ZIKV-specific binding antibody levels of cohort C animals expressed as log_10_ 50% effective dilution of binding in an ELISA assay before (open bars) and 28 days after ZIKV infection (filled bars).

#### Clinical signs consistent with symptomatic ZIKV infection were not observed during secondary ZIKV infections in MCM

Mean serum ALT, CR, CPK, and AST qualitatively increased relative to baseline in cohort C animals during the first 4-10 days of infection, after which values began decreasing, some reaching levels below baseline by 28 dpi **(S9 Fig B, C, D, E)**. Although individual variations in parameters beyond the WNPRC reference ranges were observed, these differences were not determined to be clinically significant and overall, serum chemistry panels showed no significant differences between serotype exposure groups **(S9 Fig, https://go.wisc.edu/1u3nu1)**. CBC tests also showed no significant differences overall between DENV serotype exposure groups based on calculation of the AUC for each animal’s parameters after normalizing by calculating the fold change from baseline and then comparing between groups using pairwise comparisons **(S10 Fig, https://go.wisc.edu/1u3nu1)**. All groups showed transient mean decreases from baseline in HB, HCT, and PLT counts post-infection **(S10 Fig A, B, C)**. Values began returning to mean baseline levels by 28 dpi. Body temperature and body weight were recorded longitudinally throughout the study period **(S10 Fig E, F)**. No animals exhibited clinically significant fever although some individual temperatures registered above or below WNPRC reference ranges for cynomolgus macaques. Additionally, most animals experienced minor weight loss after ZIKV infection, but we could not rule out the impact of frequent sedation and sample collection on the animals. No significant difference was observed when repeated measures ANOVA was used to examine body temperature differences between serotype exposure groups over time (F(3,11) = 2.98, p = 0.078). Similarly, no significant differences were observed when comparing body weight over the course of the study between serotype exposure groups (F(3,1) = 0.108, p = 0.74) using repeated measures ANOVA.

## Discussion

The antigenic similarities between DENV and ZIKV have led to concerns that prior infection with, or vaccination against, one virus impacts the severity of disease upon secondary infection with the other virus (48). Several lines of laboratory evidence support the possibility that DENV-specific antibodies can enhance ZIKV replication, serving as the impetus for the current study. Here, we directly compare the influence of all four DENV serotypes on subsequent ZIKV disease. The presence of heterotypic binding but non-neutralizing antibodies in human secondary DENV infections can be associated with increased viral load and disease severity (41, 49). However, within the context of secondary ZIKV infection, we did not observe a difference in either the magnitude or duration of ZIKV vRNA detection in the plasma of cohort A animals compared with ZIKV control animals (Fig 3). Consistent with these data, there were no significant differences in clinical laboratory parameters, body weight, or temperature between cohort A and negative control animals, although in some instances, we did observe differences compared with ZIKV control animals. Overall, the laboratory parameters suggest that the frequency of blood draws and animal manipulation, every day for the first 10 days of the study, may explain transient chemistry panel and CBC test perturbations. Alternatively, we may have observed natural fluctuations in parameters on an individual level that were only detectable due to the high frequency of sampling. This observation highlights the importance of including negative control animals for data interpretation. All cohort A animals had clinically in-apparent ZIKV infections, consistent with historical data for ZIKV infections in our animals (24, 50). Taken together, these results suggest that within a year of DENV-3 infection there is no evidence of protection from nor enhancement of ZIKV infection in rhesus macaques. These findings are consistent with those reported by Pantoja et al. and by McCracken et al. for rhesus macaques infected with ZIKV after DENV-1, DENV-2, DENV-4, or yellow fever virus (YFV) infection (30, 51).

Cohort B animals allowed us to examine the influence of primary ZIKV infection on secondary DENV infection. We found that the peak DENV-2 plasma vRNA loads of cohort B animals were higher than the DENV-2 serum vRNA loads for four DENV control animals, though this is likely due to the different sample types tested for vRNA in the two groups (Fig 5). There were no clinical signs of severe DENV infection nor were any of the clinical and laboratory parameters perturbed in a way that suggested enhanced disease in any of the animals. The presence of heterotypic Ab is associated with increased DENV disease in secondary DENV infections (52). Interestingly, prior to DENV-2 infection, we detected both binding and a low level of nAb in the serum of cohort B animals. This in contrast to cohort A animals for whom we did not detect ZIKV nAb after DENV infection and prior to ZIKV infection. Despite this, we observed no outward signs of DENV disease nor ADE in cohort B animals based on the clinical and laboratory parameters we examined. Overall, our results show that although DENV-2 bAb and nAb are present prior to DENV-2 infection, prior ZIKV exposure does not confer protection from infection or enhancement of DENV-2 disease in rhesus macaques. This may be different in animals exposed to DENV-2 many years after ZIKV infection. Our findings seemingly contradict those reported by George et al. who observed elevated body temperature, neutropenia, lymphocytosis, and hyperglycemia, as well as significantly enhanced peak DENV-2 plasma viremia, in rhesus macaques infected with DENV after primary ZIKV exposure (42).

To examine whether pre-existing immunity to any of the four DENV serotypes and additionally, to model a natural exposure route within the context of secondary ZIKV infection, we compared ZIKV infections across four groups of MCM. Each group was previously exposed to a different serotype of DENV by SC inoculation at a consistent time point. Even though ZIKV-bAb were present in the plasma of all cohort C animals and nAb were detected in the serum of cohort C-1 (previously infected with DENV-1), we observed no evidence of protection from, nor enhancement of, ZIKV disease in any animals. One notable limitation of our findings for cohort C is the lack of ZIKV-alone or mock-inoculated control animals of the same species. The MCM with prior DENV exposure were available opportunistically for ZIKV infection, but DENV-naive MCM were not available to use as simultaneous control animals. While this may limit our ability to directly compare ZIKV infections in DENV-exposed with DENV-naive animals, we are still able to examine whether different DENV serotype exposures differentially influence ZIKV outcome. Because DENV-3 exposure in rhesus macaques did not result in protection or enhancement of ZIKV disease relative to DENV-naive control animals, and there was no difference in ZIKV outcome between DENV-3 and any other DENV serotype in MCMs, we postulate that other DENV serotypes likely also do not alter ZIKV disease relative to DENV-naive animals. In addition, the vRNA loads presented here were consistent in both magnitude and duration with previously published studies of ZIKV infection in cynomolgus macaques (26, 27). Likewise, the delay in the time to peak ZIKV RNA loads observed for the mosquito-infected animals in cohort C compared with SC-inoculated animals is consistent with those described previously for rhesus macaques infected with ZIKV via mosquito bite (47). Interestingly, when we compared the AUC for each cohort C animal’s vRNA load, mosquito-infected animals had higher AUC values than SC-inoculated animals, suggesting that the mosquito infections may have resulted in higher vRNA loads and/or a longer duration of vRNA detection in these animals. We previously observed a similar trend with 2 of 4 rhesus macaques infected with ZIKV via mosquito bite having peak ZIKV RNA loads approximately 0.5-1 log_10_ higher than SC-inoculated macaques (47). Potentially, the difference in magnitude could be the result of mosquito-infected animals receiving multiple infectious bites. Cohort C1 had detectable ZIKV nAb prior to infection; however, we did not identify statistically significant differences in plasma vRNA loads between groups based on DENV serotype exposure history (Fig 6). Likewise, there was no consistent viral load pattern based on the estimated number of mosquito bites/probes each animal received (**S1 Table**).

Overall, our studies of secondary ZIKV infection within a year of primary DENV infection in macaques suggest that there is no effect of pre-existing DENV immunity on ZIKV infection in macaque monkeys, consistent with other macaque studies (30, 51). This is in contrast with in vitro and murine studies that have shown significant enhancement of secondary ZIKV infection in the presence of anti-flavivirus antibodies (12, 15, 53, 54). Whether this is consistent with secondary ZIKV infections in humans is unknown, but recent human cohort studies suggest more similarities with macaques (5, 21, 22, 54). In addition, there was, at most, a minimal difference in peak DENV vRNA load between animals with and without prior ZIKV exposure, suggesting that prior ZIKV exposure may only minimally affect DENV disease, at least in macaques. Primary and secondary DENV infections in macaques are largely subclinical while in humans, approximately 25% of DENV infections are estimated to be symptomatic (55). Likewise, based on a meta-analysis that included 23 epidemiological studies by Haby et al. in June of 2018, approximately 60% of human ZIKV infections are estimated to be symptomatic while most macaques show no signs of infection (24–30, 56). This suggests that findings in macaques may not entirely recapitulate disease observed in humans for either DENV, ZIKV, or sequential infections with both. However, macaques remain a relevant model for human disease, in particular, in pregnancy studies where no other animal model as closely mimics human pregnancy and congenital Zika syndrome (57–62).

Although we found no evidence of enhanced ZIKV infection in DENV-immune macaque monkeys, it is important to note that this was in nonpregnant animals. These data therefore do not address the potential impact of pre-existing DENV immunity on ZIKV infection during pregnancy. Pregnancy is associated with major immunological changes that are likely related to the maintenance of an allogeneic fetus, including changes in the systemic cytokine milieu, impaired B cell lymphopoiesis, and high numbers of tolerogenic and regulatory T and B cells (63–66). Thus, it will be imperative to determine if pregnancy-specific cofactors impact potential interactions between ZIKV pathogenesis and DENV immunity.

## Materials and Methods

### Ethics Statement

The macaques used in this study were cared for by the staff at the Wisconsin National Primate Research Center (WNPRC) in accordance with recommendations of the Weatherall report and the principles described in the National Research Council's Guide for the Care and Use of Laboratory Animals (67). The University of Wisconsin - Madison, College of Letters and Science and Vice Chancellor for Research and Graduate Education Centers Institutional Animal Care and Use Committee approved the nonhuman primate research covered under protocol number G005401-R01. The University of Wisconsin - Madison Institutional Biosafety Committee approved this work under protocol number B00000117. The use of mice to infect mosquitoes with ZIKV in this study was approved by the University of Wisconsin-Madison, School of Veterinary Medicine Institutional Animal Care and Use Committee under protocol number V005519. Mice were housed at the University of Wisconsin-Madison Mouse Breeding Core within the School of Medicine and Public Health. Once infected with ZIKV, they were housed in the Department of Pathobiological Sciences BSL-3 Insectary facility.

All animals were housed in enclosures with required floor space and fed using a nutritional plan based on recommendations published by the National Research Council. Animals were fed a fixed formula, extruded dry diet with adequate carbohydrate, energy, fat, fiber, mineral, protein, and vitamin content. Macaque dry diets were supplemented with fruits, vegetables, and other edible objects (e.g., nuts, cereals, seed mixtures, yogurt, peanut butter, popcorn, marshmallows, etc.) to provide variety to the diet and to inspire species-specific behaviors such as foraging. To further promote psychological well-being, animals were provided with food enrichment, structural enrichment, and/or manipulanda. Environmental enrichment objects were selected to minimize chances of pathogen transmission from one animal to another and from animals to care staff. While on study, all animals were evaluated by trained animal care staff at least twice each day for signs of pain, distress, and illness by observing appetite, stool quality, activity level, physical condition. Animals exhibiting abnormal presentation for any of these clinical parameters were provided appropriate care by attending veterinarians. Prior to all minor/brief experimental procedures, macaques were sedated using ketamine anesthesia and monitored regularly until fully recovered from anesthesia. Mice were anesthetized using isoflurane prior to inoculation and CO2 was used as the euthanasia method.

### Macaques

Nine male and four female Indian-origin rhesus macaques (*Macaca mulatta)* and fifteen male Mauritian cynomolgus macaques (*Macaca fascicularis)* comprising the experimental cohorts utilized in these studies were housed and cared for at the WNPRC. Animals were observed daily and samples including blood, body weight, and body temperature measurements were collected as described previously with a timeline as shown in Fig 1 (47). Historical data for ZIKV control and DENV control rhesus macaques were collected for previous, unrelated studies.

### Viruses

ZIKV strains used in these studies included: Zika virus/H.sapiens-tc/FRA/2013/FrenchPolynesia-01_v1c1 (ZIKV-FP) and Zika virus/H.sapiens-tc/PUR/2015/PRVABC59-v3c2 (ZIKV-PR). ZIKV-FP was originally obtained from Xavier de Lamballerie (European Virus Archive, Marseille, France). ZIKV-PR was obtained from Brandy Russell (CDC, Ft. Collins, CO). Both ZIKV strains were prepared as described previously (24, 47). The DENV-3 strain used to infect cohort A animals was dengue virus/H.sapiens-tc/IDN/1978/Sleman/78 (DENV-3 throughout the text). The DENV-2 strain used to infect cohort B was dengue virus/H.sapiens-tc/NGU/1944/NGC-00982_p17c2 (NGC) and was obtained from Brandy Russell (CDC, Ft. Collins, CO) (DENV-2 throughout the text). The four DENV strains used to infect the MCM (cohort C) include: DENV-1, dengue virus/H.sapiens-tc/NRU/1974/WP74, DENV-2 (NGC as above), DENV-3 (Sleman/78 as above), and DENV-4, dengue virus/H.sapiens-tc/IDN/1978/1228. DENV-1 and DENV-3 were originally obtained from the NIH while DENV-2 and DENV-4 were obtained from the CDC (Ft. Collins, CO). All four viruses for cohort C were prepared by Takeda Vaccines, Inc. (Cambridge, MA).

### Animal infections

Cohort A animals (n=3 rhesus macaques) were infected subcutaneously (SC) with 0.5 mL of 6 × 10^5^ PFU/0.5mL of DENV-3. Approximately 6-12 months later, they were SC-inoculated with 1 mL of 1 × 10^4^ PFU/mL of ZIKV-FP (24, 34) (Table 1 **and S1 Table**). cohort B animals (n=3) were SC-inoculated twice, 70 days apart, with 1 mL of 1 × 10^4^ PFU/mL ZIKV-FP and approximately one year later they were infected SC with 1 mL of 1 × 10^5^ PFU/mL of DENV-2 (Table 1 and **S1 Table**). Fifteen Mauritian cynomolgus macaques (MCM, cohort C) were SC-inoculated with 0.5 mL of 1 × 10^5^ PFU/0.5 mL DENV and approximately one year later with ZIKV-PR by mosquito-bite (n=13), or a 1 mL SC inoculation (n=2, 1 × 10^4^ PFU/mL) (Table 1 **and S1 Table**). Rhesus macaque negative control animals (n=3) were SC-inoculated with 1 mL sterile PBS. Rhesus macaque ZIKV control animals were previously SC-inoculated with ZIKV (10^4^ PFU/mL) for other studies (24, 34, 50). DENV control animal serum vRNA data for cohort B animals were obtained from Takeda Vaccines, Inc. and included five rhesus macaques infected via SC inoculation with 1 × 10^5^ PFU/0.5mL DENV-2. Detailed descriptions of each cohort can be found in **S1 Table** and the sampling timeline after each infection is shown in Fig 1.

### Mosquito infections

*Aedes aegypti* (black-eyed Liverpool (LVP) strain) used in this study were obtained from Lyric Bartholomay (University of Wisconsin-Madison, Madison, WI) and maintained at the University of Wisconsin-Madison as previously described (68). *Ae. aegypti* LVP are ZIKV transmission competent (47, 69). Mosquitoes were exposed to ZIKV by feeding on isoflurane anesthetized, ZIKV-infected *Ifnar1-/-* mice as described previously (47). These mice yielded an average infectious blood meal concentration of 1.45 × 10^6^ PFU/mL (± 0.218, n=4). Blood-fed mosquitoes were maintained as described previously (47) and randomized prior to exposure to two groups of ZIKV-naive, anesthetized MCM (cohort C). ZIKV saliva titers collected from blood fed mosquitoes ranged from 10^1.48^ to 10^3.26^ PFU **(S8 Fig)**.

### Hematology

Complete blood count (CBC) tests were assessed from EDTA-treated whole blood using a Sysmex XS-1000i hematology analyzer and manual slide evaluations as described previously (58). Hemoglobin (HB), hematocrit (HCT), platelet (PLT), and white blood cell (WBC) counts were compared between cohort A and ZIKV and negative controls, cohort B and negative controls, and cohort C DENV serotype exposure groups. Serum chemistry panels were evaluated using a Cobas 6000 analyzer (Roche Diagnostics, North America). Serum aspartate aminotransferase (AST), alanine aminotransferase (ALT), creatinine (CR), alkaline phosphatase (ALP), creatine phosphokinase (CPK), and lactate dehydrogenase (LDH) levels were compared between groups as described for CBC test parameters.

### Plaque reduction neutralization test (PRNT_50_)

Titers of ZIKV or DENV neutralizing antibodies were determined using plaque reduction neutralization tests (PRNT) on Vero cells (ATCC #CCL-81) with a cutoff value of 50% (PRNT_50_) (70). Neutralization curves were generated in GraphPad Prism (San Diego, CA) and the resulting data were analyzed by nonlinear regression to estimate the dilution of serum required to inhibit 50% Vero cell culture infection.

### Dengue reporter virus particle assay

DENV-specific neutralizing antibodies were quantified in the serum of cohort C animals using a luciferase-expressing dengue reporter virus particle (RVP) neutralization assay for all four serotypes of DENV. Serum samples and positive controls were heat inactivated at 56°C for 30 minutes and diluted four-fold in Opti-MEM media (Thermo Fisher Scientific, Inc., Waltham, MA). Assays were conducted in triplicate 384-well plates using human lymphoblastoid cell line (Assay-Ready Frozen Instant Cells of Raji-DC-SIGNR; Raji cells, acCELLerate GmbH, Hillsborough, NJ) expressing flavivirus attachment factor lectin DC-SIGNR (CD209L). Plates containing diluted serum and dengue RVP were incubated at 37°C, 5% CO_2_ for 1 hour to allow formation of immune-complexes to reach equilibrium. Thereafter, 15µL of Raji-R cells diluted in Opti-MEM at 4×10^5^cells/mL seeding density were added to all wells of the plates. Plates were then incubated at 37°C, 5% CO_2_ for 72 ± 2 hours. Following incubation, plates were equilibrated to room temperature for 15 minutes followed by addition of 30 µL of Renilla-Glo (Promega Co., Madison, WI) detection reagent (diluted 1:100 in Renilla-Glo buffer). After 15 minutes, the plates were read using the Perkin Elmer EnSpire Luminescence program (Perkin-Elmer, Inc., Waltham, MA). Raw data were transposed in Microsoft Excel (Microsoft Co., Redmond, WA) to fit the format requirements of GraphPad Preference intervalSM (GraphPad Software, San Diego, CA). The titer of each sample was determined by calculating EC50 values using sigmoidal dose response nonlinear regression analysis.

### ZIKV RNA isolation and quantitative reverse transcription PCR (qRT-PCR)

Plasma and PBMC were isolated from EDTA-treated whole blood on Ficoll paque at 1860 x rcf for 30 minutes as described in Dudley et. al. (24). Serum was isolated from clot-activator tubes without additive. Viral RNA (vRNA) was extracted as previously described with a Maxwell 16 MDx instrument (Promega, Madison, WI) and evaluated using qRT-PCR (24, 34, 50). RNA concentration was determined by interpolation onto an internal standard curve of seven ten-fold serial dilutions of a synthetic ZIKV RNA segment based on ZIKV-FP. The limit of detection of this assay is 100 copies vRNA/ml plasma or serum.

### DENV quantitative reverse transcription PCR (qRT-PCR)

For DENV challenged animals (cohort B and DENV controls), vRNA in serum or plasma samples was measured using qRT-PCR at Takeda Vaccines. Viral RNA was extracted from 140µl of each sample using the QIAamp viral RNA kit (Qiagen, Valencia, CA). The vRNA was eluted in 60 µl elution buffer and stored at −80°C until use. Viral RNA was quantified in a singleplex qRT-PCR with a primer/probe set targeting the 3’ non-coding region of DENV using a standard curve derived from in vitro transcribed cDNA clones and quantified as previously described (71). All qRT-PCR reactions were performed in a final volume of 25 µl using the ABI 4X TaqMan Fast Virus 1-Step Master Mix. The reactions contained 5 µl extracted vRNA, 0.4 µM of each primer, and 0.2 µM probe. The reaction was conducted in the ABI 7500DX using a cycling protocol as follows: cycle 1 - 50°C for 5 minutes, cycle 2 - 95°C for 20 seconds, repeat cycle 3, 45 times - 95°C for 3 seconds and 55°C for 30 seconds. The qRT-PCR limit of detection of 3.6 log_10_ copies vRNA/mL was determined by testing nine replicates per dilution of the standard curve and selecting the concentration with a 100% detection rate as well as a low (≤ 0.5) cycle threshold standard deviation of the replicates.

### ZIKV and DENV whole-virion binding ELISA

High-binding 96-well ELISA plates (Greiner) were coated with 30ng 4G2 antibody (clone D1-4G2-4-15) in carbonate buffer (pH 9.6) overnight at 4°C. Plates were blocked in Tris-buffered saline containing 0.05% Tween-20 and 5% normal goat serum (cat.# G6767, Sigma-Aldrich, St. Louis, MO) for 1 hour at 37°C, followed by incubation with either ZIKV (PRVABC59, BEI) or DENV-2 (New Guinea C, BEI) for 1 hour at 37°C. Heat inactivated plasma was tested at an 1:12.5 starting dilution in 8 serial 4-fold dilutions in duplicate, incubating for 1 hour at 37°C. Horseradish peroxidase (HRP)-conjugated goat anti-monkey IgG antibody (Abcam, Cambridge, MA) was used at a 1:2,500 dilution, followed by the addition of SureBlue reserve TMB substrate (KPL, Gaithersburg, MD). Reactions were stopped by stop solution (KPL, Gaithersburg, MD). Optical densities (OD) were detected at 450 nm. The limit of detection was defined as an OD value of the 1:12.5 dilution greater than three times the background OD of ZIKV/DENV naive macaque serum. The log_10_ 50% effective dilutions (ED_50_) were calculated for IgG binding responses against the whole virion and compared between 0 and 28 dpi time points.

### Cytokine and Chemokine Profiling Using Luminex

Plasma samples from cohort A rhesus macaques were analyzed using Milliplex map Nonhuman Primate Cytokine Magnetic Bead Panel Premixed 23-Plex Assay (EMD Millipore Corporation, Billerica, MA). Assays were run on a Bio-Plex 200 system and analyzed using Bio-Plex Manager Software version 6.1.1 (Bio-Rad Laboratories, Hercules, CA). Standard curves were calculated using a logistic-5PL regression method using Bio-Plex Manager software version 6.1.1 (Bio-Rad Laboratories, Hercules, CA).The following 23 cytokines and chemokines are included in the panel: G-CSF, GM-CSF, IFN-γ, IL-1ra, IL-1β, IL-2, IL-4, IL-5, IL-6, IL-8, IL-10, IL-12/23 (p40), IL-13, IL-15, IL-17, IL-18, MCP-1 (CCL2), MIP-1α (CCL3), MIP-1β (CCL4), sCD40L, TGF-α, TNF-α, and VEG. An additional dilution of the standard, beyond what is suggested in the manufacturer’s protocol, was included and used in analysis when detectable above background. Samples from animals 489988, 756591, 875914, 850585, 411359, and 321142 were assessed using an 8-point standard curve. 448436, 774011, and 829256 were assessed using a 7-point standard curve. With the exception of the additional dilution of the standard, the assay was performed according to the manufacturer's protocol and used the provided serum matrix as a background control. To minimize plate effect as a confounder in analyses, all plasma samples from individual animals were assayed on a single plate. With the exceptions of IL-1ra, IL-8, IL-2, IL-15, MCP-1 (CCL-2), and sCD40L, the cytokine and chemokine levels were not discernible above background levels of fluorescence and were not interpretable.

### Body Weight and Temperature Measurements

Body weight and body temperature measurements were collected as shown in Fig 2 to assess weight loss and fever as proxies for disease. Body weights were monitored by WNPRC animal care and veterinary staff throughout the studies. Body weight data were not collected for cohort B animals. Temperatures were compared with the WNPRC reference ranges for the appropriate macaque species when determining the presence and/or absence of fever at each time point. WNPRC veterinary staff were consulted in determining whether an individual animal’s body temperature outside the reference ranges was clinically significant.

### Statistical Analyses

Longitudinal vRNA loads (vRNA/mL plasma or serum) were compared between ZIKV control animals and cohort A, between DENV control animals and cohort B, and between cohorts C1-C4 over time, using Student’s t-tests (cohorts A and B) or analysis of variance (ANOVA) (cohort C) after calculating the area under the curve (AUC) for each animal’s vRNA load trajectory in R Studio (v.1.1.383). Peak vRNA loads were compared between ZIKV control animals and cohort A, between DENV control animals and cohort B, and between C1-C4 using Student’s t-tests or non-parametric equivalents (cohorts A and B) or ANOVA (cohort C) in R Studio (v.1.1.383).

CBC test and serum chemistry panel parameters were normalized to baseline (pre-infection) levels by calculating fold changes from the baseline. The magnitude of laboratory values over the 28 day follow-up period was quantified by calculating the area under the curve (AUC) using the trapezoidal rule for each animal and laboratory parameter based on the fold changes from day 0 to day 28. Because the AUC values were non-normally distributed, all AUC values were log-transformed before conducting the analysis. Analysis of variance (ANOVA) was used to compare the log-transformed AUC values between groups. Longitudinal changes of the fold changes were compared between groups using a linear mixed effects model with animal specific random effects. Multiple comparisons between groups were conducted using Tukey’s Honestly Significant Difference (HSD) to control the type I error. All reported p-values are two-sided and p < 0.05 was used to define statistical significance. Statistical analyses of CBC test and serum chemistry panel data were conducted using SAS software v. 9.4 (SAS Institute Inc., Cary NC).

Differences in IL-1ra, IL-8, IL-2, IL-15, MCP-1, and sCD40L values were normalized to baseline values, transformed to positive values, and compared between negative control, ZIKV control, and cohort A animals using repeated measures ANOVA followed by pairwise comparisons where appropriate using Tukey’s HSD in R Studio (v.1.1.383).

Body weights were compared between cohort A, ZIKV control, and negative control animals using a mixed effects model with weight as the dependent variable, dpi as the fixed effect, and animal ID as a random effect. Longitudinal body weights were compared between DENV serotype exposures for cohort C using two-way ANOVA in the lme4 package (72). Longitudinal body temperatures were compared between cohort A and ZIKV control and negative control animals, between cohort B and negative control animals using mixed effects models with temperature as the dependent variable, cohort/group/serotype as the fixed effect, and animal ID as the random effect using the lme4 package (72). Final mixed effects models were chosen based on the minimization of Akaike information criteria (AIC). Body temperatures were compared between cohort C DENV serotype exposure groups using repeated measures ANOVA. All body weight and temperature data were analyzed in R Studio (v.1.1.383).

### Data management

Complete datasets for these studies have been made publicly available in a manuscript-specific folder on the Zika Open Research Portal (https://go.wisc.edu/6wrw87). Authors declare that all other data for these study findings are available via this portal or through supplementary information files from this article.

## Acknowledgements

G. Young and H. Dean are employees of Takeda Vaccines, Inc. All other authors declare no conflicts of financial or personal interests. We thank the Veterinary Services, Colony Management, Scientific Protocol Implementation, and the Pathology Services staff at the Wisconsin National Primate Research Center (WNPRC) for their contributions to this study. We thank Xavier de Lamballerie and Brandy Russell for providing virus isolates. We also thank Lyric Bartholomay for providing mosquitoes for cohort C animal infections. We acknowledge Jens Kuhn and Jiro Wada for preparing the silhouettes of macaques used in figures.

